# Falciparum malaria from coastal Tanzania and Zanzibar remains highly connected despite effective control efforts on the archipelago

**DOI:** 10.1101/863019

**Authors:** Andrew P Morgan, Nicholas F Brazeau, Billy Ngasala, Lwidiko E. Mhamilawa, Madeline Denton, Mwinyi Msellem, Ulrika Morris, Dayne L Filer, Ozkan Aydemir, Jeffrey A. Bailey, Jonathan Parr, Andreas Mårtensson, Anders Bjorkman, Jonathan J Juliano

**Author notes:** Co-first authors (Andrew P Morgan, Nicholas F Brazeau). Corresponding Author: Jonathan Juliano, University of North Carolina, CB#7030, 130 Mason Farm Rd., Chapel Hill, NC 27599.

## Abstract

**Background:** Tanzania’s Zanzibar archipelago has made significant gains in malaria control over the last decade and is a target for malaria elimination. Despite consistent implementation of effective tools since 2002, elimination has not been achieved. Importation of parasites from outside of the archipelago is thought to be an important cause of malaria’s persistence, but this paradigm has not been studied using modern genetic tools.

**Methods:** We used whole-genome sequencing (WGS) to investigate the impact of importation, employing population genetic analyses of *Plasmodium falciparum* isolates from both the archipelago and mainland Tanzania. We assessed ancestry, levels of genetic diversity and differentiation, patterns of relatedness, and patterns of selection between these two populations by leveraging recent advances in deconvolution of genomes from polyclonal malaria infections.

**Results:** We identified significant decreases in the effective population sizes in both populations in the timeframe of decreasing malaria transmission in Tanzania. Identity by descent analysis showed that parasites in the two populations shared large sections of their genomes, on the order of 5 cM, suggesting shared ancestry within the last 10 generations. Even with limited sampling,, we demonstrate a pair of isolates between the mainland and Zanzibar that are related at the expected level of half-siblings, consistent with recent importation

**Conclusions:** These findings suggest that importation plays an increasing role for malaria incidence on Zanzibar and demonstrate the value of genomic approaches for identifying corridors of parasite movement to the island.

## BACKGROUND

Despite nearly two decades of progress in control, malaria remains a major public health challenge with an estimated 219 million cases and 435,000 deaths in 2017 globally [1]. The mainland of Tanzania has heterogeneous transmission of mainly *Plasmodium falciparum* malaria, but overall levels of malaria remain high, accounting for approximately 3% of global malaria cases [1]. However, through a combination of robust vector control and access to efficacious antimalarial treatment, the archipelago of Zanzibar has been deemed a pre-elimination setting, having only low and mainly seasonal transmission [2]. Despite significant efforts, however, elimination has been difficult to achieve in Zanzibar. The reasons for Zanzibar’s failure to achieve elimination are complex and likely driven by several key factors: 1) as transmission decreases, the distribution of cases changes and residual transmission is more focal and mainly outdoors [3]; 2) a significant number of malaria infections are asymptomatic and thus untreated and remain a source for local transmission [4–7]; and 3) the archipelago has a high level of connectivity with the mainland, thus imported malaria through human travel may play an increasing relative role in transmission.

Genomic epidemiology can supplement traditional epidemiological measures in studies of malaria transmission and biology, thereby helping to direct malaria elimination strategies [8]. Whole-genome sequencing (WGS) can be particularly useful for understanding the history of parasite populations and movement of closely related parasites over geographical distances [9,10]. Identity by descent (IBD), the sharing of discrete genomic segments inherited from a common genealogical ancestor, has been found to be a particularly good metric for studying the interconnectivity of parasite populations [11–13]. A major obstacle to studying IBD in microorganisms, and in particular malaria, is the presence of multiple clones in a single infection. In order to address this obstacle, recent algorithms have been developed to deconvolute multiple infections into their respective strains from Illumina sequence data [14,15]. These advances now make it tractable to conduct population genetic analysis of malaria in regions of higher transmission, where infections are often polyclonal.

Decreases in malaria prevalence are hypothesized to be associated with increasing clonality of malaria population, decreased overall parasite diversity and a reduced complexity of infection (COI), defined as a decreased number of infecting clones [8]. This has been shown in pre-elimination settings in Asia as well as in lower transmission regions of Africa [16–18]. It has not been determined if a similar reduction in diversity has occurred in Zanzibar with the significant reduction of malaria in the archipelago. We evaluated if a population contraction has occurred in the *P. falciparum* parasites from the pre-elimination region of the Zanzibar archipelago compared to parasites from mainland Tanzania using WGS data. We used the sequence data to: 1) assess the ancestry of parasites in the two regions, 2) determine the levels of genetic diversity and differentiation, 3) determine patterns of relatedness and inbreeding and 4) assess for signatures of adaptation and natural selection. We then use this information and IBD analysis to assess for genetic signatures of recent importation of parasites from the higher transmission regions of mainland Tanzania to the lower transmission regions of the Zanzibar archipelago to better understand how importation is affecting malaria elimination efforts.

## METHODS

### Clinical samples

WGS was attempted on 106 *Plasmodium falciparum* samples collected from subjects with uncomplicated malaria or asymptomatic infection from 2015 to 2017. Forty three of these were leukodepleted blood collected as part of an *in vivo* efficacy study of artemether-lumefantrine (AL) in pediatric uncomplicated malaria patients collected from 2015-2017 in Yombo, Bagamoyo District. A remaining 63 samples were dried blood spots (DBS) collected in Zanzibar in 2017. These samples came from cross-sectional surveys of asymptomatic individuals (*n* = 34) and an *in vivo* efficacy study of artesunate-amodiaquine (ASAQ) with single low dose primaquine (SLDP) in pediatric uncomplicated malaria patients (*n* = 29). The participants from Zanzibar also provided travel histories for any travel off the archipelago in the last month. Clinical characteristics of the attempted and sequenced samples from each cohort from Zanzibar is provided in **Supplemental Table 1**.

### Generation and sequencing of libraries

Leukodepleted blood samples and DBS were extracted using QIAmp 96 DNA blood kits per the manufacturer protocol (Qiagen, Hilden, Germany). DNA from leukodepleted blood was acoustically sheared using a Covaris E220 instrument, prepared for sequencing without enrichment using Kappa Hyper library preps, and individually barcoded per manufacturer’s protocol (Kappa Biosystems, Columbus, OH). DNA extracted from DBS was enriched for *P. falciparum* DNA before library prep using two separate selective whole genome amplification (sWGA) reactions. The sWGA approach was adapted from previously published methods and employed two distinct sets of primers designed for *P. falciparum*, including the Probe_10 primer set described previously by Oyola *et al.* and another set of custom primers (JP9) we designed using ‘swga’[19–21]. We included phosphorothioate bonds between the two most 3’ nucleotides for all primers in both sets to prevent primer degradation. Design and evaluation of these custom primers and the sWGA approach are described in the **Supplemental Materials** and **Supplementary Table 2**. The two sWGA reactions were carried out under the same conditions. The products of the two sWGA reactions were pooled in equal volumes and acoustically sheared using a Covaris E220 instrument before library preparation using Kappa Hyper library preps. The indexed libraries were pooled and sequenced on a HiSeq4000 using 2×150 chemistry at the University of North Carolina High Throughput Sequencing Facility. Sequencing reads were deposited into the NCBI SRA (Accession numbers: pending).

### Public sequencing data

Illumina short read WGS data for *Plasmodium falciparum* isolates was downloaded from public databases. This included 68 isolates from other regions of Tanzania, collected between 2010 and 2013, as well as 179 isolates from other regions, including Southeast Asia, South Asia, East and West Africa (**Supplemental Table 3**).

### Read alignment and quality control

Raw paired-end reads were trimmed for adapter sequences with ‘cutadapt’ v1.18 and aligned to the *P. falciparum* 3D7 reference genome (assembly version 3, PlasmoDB version 38: https://plasmodb.org/common/downloads/release-38/Pfalciparum3D7/fasta/data/PlasmoDB-38_Pfalciparum3D7_Genome.fasta) with ‘bwa mem’ v0.7.17-r1188. Duplicates were marked with ‘samblaster’ v0.1.24. We defined a position as “callable” if it was covered by >= 5 high-quality reads (MQ >= 25, BQ >= 25), and computed the proportion of callable sites in each isolate was calculated with the Genome Analysis Toolkit (GATK) ‘CallableLoci’ tool v3.8-0. Only isolates with >= 70% of the genome callable were used for further analysis.

### Variant discovery and filtering

Short sequence variants (including SNVs, indels and complex multi-nucleotide variants) were ascertained in parallel in each isolate using GATK ‘HaplotypeCaller’ v.4.0.3.0, then genotyped jointly across the entire cohort with GATK ‘GenotypeGVCFs’ according to GATK best practices. Variant discovery was limited to the core nuclear genome as defined by [22]. Putative SNVs only were filtered using the GATK Variant Quality Score Recalibration (VQSR) method. For training sets, we used: QC-passing sites from the *P. falciparum* Genetic Crosses Project release 1.0 (ftp://ngs.sanger.ac.uk/production/malaria/pf-crosses/1.0/; [22]) (true positives, prior score Q30); QC-passing sites from the Pf3K release v5.1 (ftp://ngs.sanger.ac.uk/production/pf3k/release_5/5.1/) (true positives + false positives, prior score Q15). We used site annotations QD, MQ, MQRankSum, ReadPosRankSum, FS, SOR and trained the model with 4 Gaussian components. A VQSLOD threshold −0.0350 achieved 90% sensitivity for re-discovering known sites in the training sets. All biallelic SNVs with VQSLOD at or above this threshold were retained.

Isolates may contain multiple strains that are haploid resulting in mixed infections with arbitrary effective ploidy. To account for this complexity of infection (COI) in our analyses, we followed previous authors [23] and calculated the following quantities at each variant site: for each isolate, the within-sample allele frequency (WSAF), the proportion of mapped reads carrying the non-reference allele; the population-level allele frequency (PLAF), the mean of within-sample allele frequencies; and the population-level minor allele frequency (PLMAF), the minimum of PLAF or 1-PLAF. These calculations were performed with ‘vcfdo wsaf’ (https://github.com/IDEELResearch/vcfdo).

### Analyses of mutational spectrum

Ancestral versus derived alleles at sites polymorphic in *P. falciparum* were assigned by comparison to the outgroup species *P. reichenowi*. Briefly, an approximation to the genome of the *P. reichenowi* – *P. falciparum* common ancestor (hereafter, “ancestral genome”) was created by aligning the *P. falciparum* 3D7 assembly to the *P. reichenowi* CDC strain assembly (version 3, PlasmoDB version 38: https://plasmodb.org/common/downloads/release-38/PreichenowiCDC/fasta/data/PlasmoDB-38_PreichenowiCDC_Genome.fasta) with ‘nucmer’ v3.1 using parameters “-g 500 -c 500 -l 10” as in [24]. Only segments with one-to-one alignments were retained; ancestral state at sites outside these segments was deemed ambiguous. The one-to-one segments were projected back into the 3D7 coordinate system. Under the assumption of no recurrent mutation, any site polymorphic in *P. falciparum* is not expected to also be mutated on the branch of the phylogeny leading to *P. reichenowi*. Thus, the allele observed in *P. reichenowi* is the ancestral state conditional on the site being polymorphic. Transitions-transversion (Ti:Tv) ratios and mutational spectra were tallied with ‘bcftools stats’ v1.19.

### Analyses of ancestry and population structure

VQSR-passing sites were filtered more stringently for PCA to reduce artifacts due to rare alleles and missing data. Genotype calls with GQ < 20 or DP < 5 were masked; sites with < 10% missing data and PLMAF >5% after sample-level filters were retained for PCA, which was performed with ‘akt pca’ v3905c48 [25]. For calculation of *f*_3_ statistics, genotype calls with GQ < 10 or DP < 5 were masked; sites with <10% missing data and PLMAF >1% after sample-level filters were retained. Then *f*_3_ statistics were calculated from WSAFs rather than nominal diploid genotype calls, using ‘vcfdo f3stat’.

### Estimation of sequence diversity

Estimates of sequence diversity and differentiation were obtained from the site-frequency spectrum (SFS), which in turn was estimated directly from genotype likelihoods with ‘ANGSD’ 0.921-11-g20b0655 [26] using parameters “-doCounts 1 – doSaf 1 -GL 2 -minDepthInd 3 -maxDepthInd 2000 -minMapQ 20 -baq 1 -c 50.” Unfolded SFS were obtained with the ‘ANGSD’ tool ‘realSFS’ using the previously-described ancestral sequence from *P. reichenowi*. All isolates were treated as nominally diploid for purposes of estimating the SFS because we noted systematic bias against mixed isolates when using ‘ANGSD’ in haploid mode. Four-fold degenerate and zero-fold degenerate sites were defined for protein-coding genes in the usual fashion using transcript models from PlasmoDB v38. SFS for all sites, 4-fold and 0-fold degenerate sites were estimated separately in mainland Tanzania and Zanzibar isolates in non-overlapping 100 kb bins across the core genome. Values of sequence diversity (theta_pi) and Tajima’s *D* were estimated for these bin-wise SFS using ‘sfspy summarize’ (https://github.com/IDEELResearch/sfspy), and confidence intervals obtained by nonparametric bootstrap. *F*_st_ was calculated from the joint SFS between mainland Tanzania and Zanzibar. The distribution of local *F*_st_ values was calculated in 5 kb bins for purposes of visualization only.

### Strain deconvolution and inheritance-by-descent analyses

Complexity of infection (COI) and strain deconvolution (phasing) were performed jointly using ‘dEploid’ v0.6-beta [14]. For these analyses we limited our attention to 125 isolates from mainland Tanzania and Zanzibar (57 new in this paper and 68 previously published). On the basis of the analyses shown in **Figures 1** and **2**, these isolates appeared to constitute a reasonably homogeneous population, so we used the set of 125 for determination of PLAFs to be used as priors for the phasing algorithm. Phasing was performed using population allele frequencies as priors in the absence of an external reference panel known to be well-matched for ancestry. We further limited the analysis to very high-confidence sites: VQSLOD > 8, 75% of isolates having GQ ≥ 10 and DP ≥ 5, ≥ 10 bp from the nearest indel (in the raw callset), ≥ 10 total reads supporting the non-reference allele, and PLMAF ≥ 1%. The ‘dEploid’ algorithm was run in “-noPanel” mode with isolate-specific dispersion parameters (“-c”) set to the median coverage in the core genome, and default parameters otherwise. Within-isolate IBD segments were extracted from the ‘dEploid’ HMM decodings by identifying runs of sites with probability ≥ 0.90 assigned to hidden states where at least two of the deconvoluted haplotypes were IBD. The total proportion of strain genomes shared IBD (within-isolate *F*_IBD_) for isolates with COI > 1 was obtained directly from ‘dEploid’ log files, and agreed closely with the sum of within-isolate IBD segment lengths.

**Figure 1.**
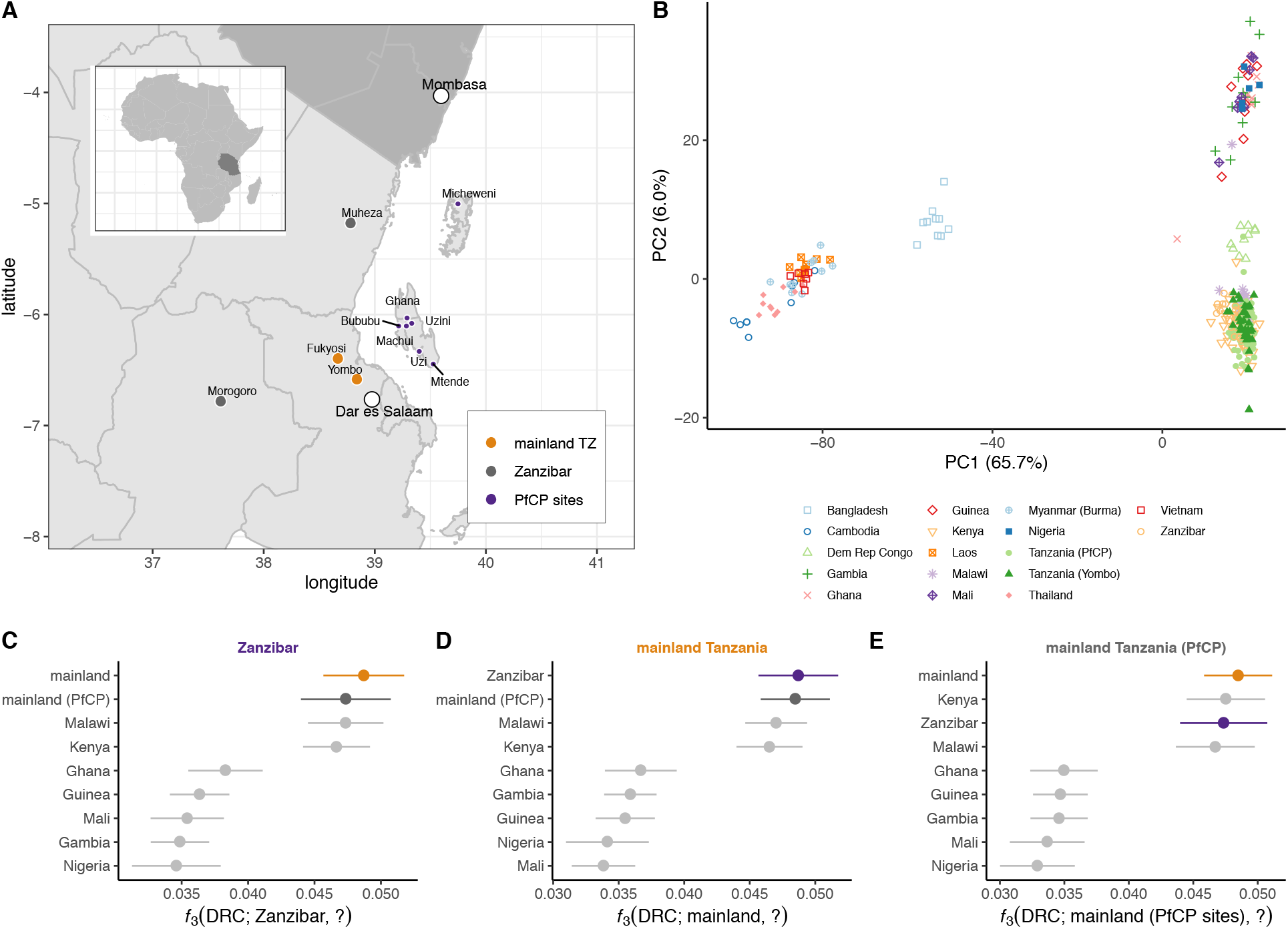
Ancestry of *P. falciparum* in Zanzibar and mainland Tanzania. (**A**) Location for samples used in this study, colored by population: orange, mainland Tanzania; purple, Zanzibar; dark grey, published mainland Tanzania isolates from the *P. falciparum* Community Project. Other major regional cities show with open circles. (**B**) Major axes of genetic differentiation between global *P. falciparum* populations demonstrated by principal components analysis (PCA) on genotypes at 7,122 SNVs with PLMAF > 5%. Each point represents a single isolate (*n* = 304) projected onto the top two principal components (71% cumulative variance explained); color-shape combinations indicate country of origin. (**C-E**) Population relationships assessed by *f*_3_ statistics with focal population indicated at the top of each panel, comparator populations on the vertical axis, and Congolese population as an outgroup. Error bars show 3 times the standard error computed by block-jacknife.

**Figure 2.**
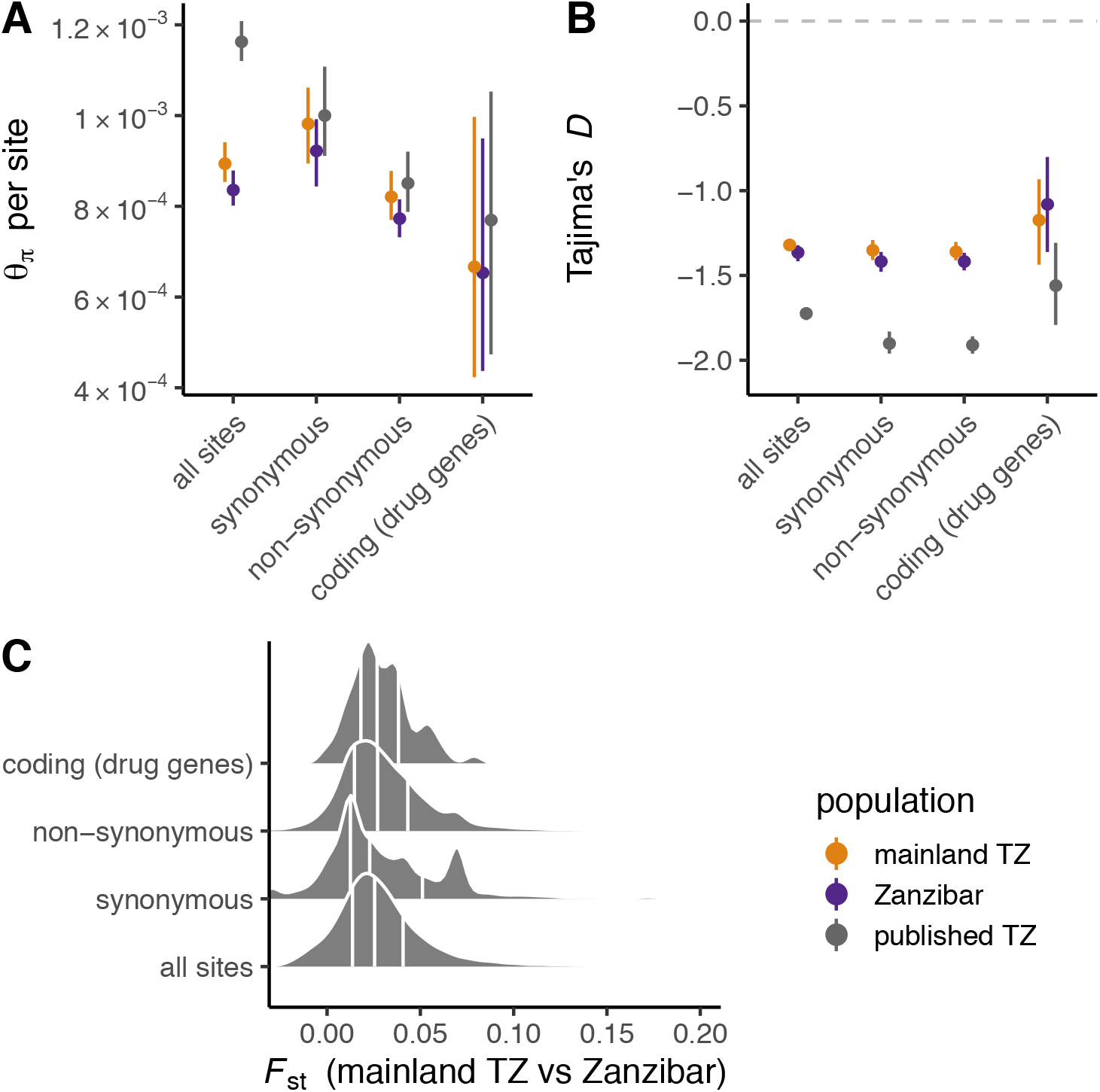
Diversity and differentiation of *P. falciparum* in mainland Tanzania and Zanzibar. (**A**) Average pairwise sequence diversity (theta_pi) per base pair in different compartments of the core genome: all sites, 4-fold degenerate (“synonymous”) sites, 0-fold degenerate (“non-synonymous”) sites, and coding regions of putative drug-resistance genes. Points are colored by population; error bars give 95% bootstrap CIs. (**B**) Tajima’s *D* in same classes of sites as in panel A. (**C**) Distribution of *F*_st_ between mainland Tanzania and Zanzibar isolates, calculated in 5 kb windows. Vertical lines mark 25th, 50th and 75th percentiles.

Between-isolate IBD segments were identified by applying ‘refinedIBD’ v12Jul18 [27] to the phased haplotypes produced by ‘dEploid’. For a genetic map, we assumed constant recombination rate of 6.44 × 10^−5^ cM/bp (equal to the total genetic length of the P. falciparum map divided by the physical size of the autosomes in the 3D7 assembly.) Segments >2 cM were retaiend for analysis. The proportion of the genome shared IBD between phased haplotypes (between-isolate *F*_IBD_) was estimated by maximum likelihood described in [28] using ‘vcfdo ibd’.

### Demographic inference

Curves of recent historical effective population size were estimated from between-isolate IBD segments with ‘IBDNe’ v07May18-6a4 [29] using length threshold > 3 cM, 20 bootstrap replicates and default parameters otherwise. Local age-adjusted parasite prevalence point estimates (*Pf*PR_2-10_) and credible intervals were obtained from the Malaria Atlas Project [30] via the R package ‘malariaAtlas’ [31].

More remote population-size histories were estimated with ‘smc++’ v1.15.2 (Terhorst et al. 2017). Phased haplotypes from ‘dEploid’ were randomly combined into diploids and parameters estimated separately for mainland Tanzania and Zanzibar populations using 5-fold cross-validation via command ‘smc++ cv’, with mutation rate set to 10^−9^ bp^−1^ gen^−1^. Marginal histories from each population were then used to estimate split times using ‘smc++ split’.

### Analyses of natural selection

The distribution of fitness effects (DFE) was estimated within mainland Tanzania and Zanzibar populations with ‘polyDFE’ v2.0 using 4-fold degenerate sites as putatively-neutral and 0-fold degenerate sites as putatively-selected [32]. “Model C” in ‘polyDFE’ parlance -- a mixture of a gamma distribution on selection coefficients of deleterious mutations and an exponential distribution for beneficial mutations -- was chosen because it does not require *a priori* definition of discrete bins for selection coefficients, and the gamma distribution can accommodate a broad range of shapes for the DFE of deleterious mutations (expected to represent the bulk of polymorphic sites.) Confidence intervals for model parameters were obtained by non-parametric bootstrap via 20 rounds of resampling over the 100 kb blocks of the input SFS. Because ‘polyDFE’ fits nuisance parameters for each bin in the SFS, we found that computation time increased and numerical stability decreased for SFS with larger sample sizes. We therefore smoothed and rescaled input SFS to fixed sample size of 10 chromosomes each using an empirical-Bayes-like method (https://github.com/CartwrightLab/SoFoS/) re-implemented in ‘sfspy smooth’. Smoothing of input SFS had very modest qualitative effect on the resulting DFE.

The cross-population extended haplotype homozygosity (XP-EHH) statistic was used to identify candidate loci for local adaptation in mainland Tanzania or Zanzibar. Because the statistic requires phased haplotypes and is potentially sensitive to phase-switch errors, only isolates with COI = 1 were used (*n* = 18 mainland Tanzania, *n* = 12 Zanzibar.) XP-EHH was calculated from haploid genotypes at a subset of 103,982 biallelic SNVs polymorphic among monoclonal isolates with the ‘xpehhbin’ utility of ‘hapbin’ v1.3.0-12-gdb383ad [33]. Raw values were standardized to have zero mean and unit variance; the resulting z-scores are known to have an approximately normal distribution [34] so nominal *p*-values were assigned from the standard normal distribution. The Benjamini-Hochberg method was used to adjust nominal *p*-values for multiple testing.

Pipelines used for WGS read alignment, variant calling, variant filtering, haplotype deconvolution and SFS estimation are available on Github: https://github.com/IDEELResearch/NGS_Align_QC_Pipelines.

## RESULTS

### WGS and variant discovery

Genomic data for *P. falciparum* was generated using leukodepleted blood collected from 43 subjects from Yombo, Tanzania (“mainland”) and from DBS collected from 63 subjects from the Zanzibar archipelago (“Zanzibar”; **Figure 1A**) using selective whole-genome amplification (sWGA) followed by Illumina sequencing. Thirty-six isolates (84%) from the mainland and 21 isolates (33%) from Zanzibar yielded sufficient data for analysis. We combined these 57 genomes with an additional 68 published genomes from other sites in Tanzania in the Pf Community Project (PfCP) and 179 genomes from other sites in Africa and Asia, representing a broad geographic sampling of Africa and Asia [35]. Single-nucleotide variants (SNVs) were ascertained jointly in the global cohort. After stringent quality control on 1.3 million putative variant sites, a total of 387,646 biallelic SNVs in the “core genome” -- the 20.7 Mb of the 3D7 reference assembly lying outside hypervariable regions and accessible by short-read sequencing [22] -- were retained for further analysis. The frequency spectrum was dominated by rare alleles: 151,664 alleles (39.1%) were singletons and 310,951 (80.2%) were present in <1% of isolates in our dataset. Ancestral and derived states at 361,049 sites (93.1%) were assigned by comparison to the *P. reichenowi* (CDC strain) genome. We observed similar biases in the mutational spectrum as have been estimated directly from mutation-accumulation experiments [36]: transitions are more common transversions (Ti:Tv = 1.12; previous estimate 1.13), with a large excess of G:C > A:T changes even after normalizing for sequence composition (**Supplementary Figure 1**). Consistency in the mutational spectrum between independent studies, using different methods for sample preparation and bioinformatics, supports the accuracy of our genotypes.

### Ancestry of mainland Tanzania and Zanzibar isolates

In order to place our isolates in the context of global genetic variation in *P. falciparum*, we used principal components analysis (PCA) (**Figure 1B**). A subset of 7,122 stringently-filtered sites with PLMAF > 5% (see **Methods**) were retained for PCA to minimize distortion of axes of genetic variation by rare alleles or missing data. Consistent with existing literature, isolates separated into three broad clusters corresponding to southeast Asia, east Africa and west Africa. Mainland Tanzania and Zanzibar isolates fell in the east Africa cluster. We formalized this observation using *f*_3_ statistics [37,38], which measure shared genetic drift in a pair of focal populations *A* and *B* relative to an outgroup population *O*. The new isolates from Yombo and Zanzibar and published Tanzanian isolates shared mutually greater genetic affinity for each other than for other populations in the panel (**Figure 1C-E**); isolates from neighboring countries Malawi and Kenya were next-closest. Together these analyses support an east African origin for parasites in mainland Tanzania and in Zanzibar.

### Genetic diversity and differentiation

In order to better understand the population demography and effects of natural selection in the parasite populations, we evaluated indices of genetic diversity within populations, and the degree to which that diversity is shared across populations. We derived several estimators of sequence diversity from the site frequency spectrum (see **Methods**) in four sequence classes: all-sites in the core genome; 4-fold degenerate (“synonymous”) sites; 0-fold degenerate (“nonsynonymous”) sites; and coding sites in genes associated with resistance to antimalarial drugs. Levels of sequence diversity were very similar within mainland Tanzania and Zanzibar isolates (theta_pi = 9.0 × 10^−4^ [95% CI 8.6 × 10^−4^ -- 9.4 × 10^−4^] vs 8.4 [95% CI 8.0 × 10^−4^ -- 8.7 × 10^−4^ per site) and 1.3-fold lower than among previously-published Tanzanian isolates (**Figure 2A**). As expected, diversity was greater at synonymous than non-synonymous sites. Tajima’s *D* took negative values in all three populations and across all sites classes (**Figure 2B**). Demographic explanations for this pattern are investigated later in the manuscript. When we evaluated differentiation between populations, we found minimal evidence for genetic differentiation between parasites in mainland Tanzania and Zanzibar.

Genome-wide *F*_st_ was just 0.0289 (95% bootstrap CI 0.0280 -- 0.0297); the distribution of *F*_st_ in 5 kb windows is shown in **Figure 2C**. These measures of between population differentiation provide minimal evidence for genetic differentiation between parasites in mainland Tanzania and Zanzibar.

### Patterns of relatedness and inbreeding

In contrast to *F*_st_, long IBD segments provide a more powerful and fine-grained view of relationships in the recent past. We took advantage of recent methodological innovations [14] to estimate complexity of infection (COI) -- the number of distinct parasite strains in a single infection -- and simultaneously obtained phased haplotypes for each strain. IBD segments were ascertained both between and (in the case of mixed infections) within isolates. We also calculated the *F*_ws_ statistic, an index of within-host diversity that is conceptually similar to traditional inbreeding coefficients [23]. Approximately half of isolates had COI = 1 (“clonal”) and half had COI > 1 (“polyclonal” or “mixed”) in both populations, and the distribution of COI was similar between the mainland and Zanzibar (chi squared = 0.27 on 2 df, *p* = 0.87; **Supplemental Table 4**). Ordinal trends in *F*_ws_ were qualitatively consistent with COI but show marked variation for COI > 1 (**Figure 3A**). We found evidence for substantial relatedness between infecting lineages within mixed isolates (**Figure 3B**): the median fraction of the genome shared IBD (*F*_IBD_) within isolates was 0.22 among mainland and 0.24 among Zanzibar isolates, with no significant difference between populations (Wilcoxon rank-sum test, *p* = 0.19). The expected sharing is 0.50 for full siblings and 0.25 for half-siblings with unrelated parents [39]. We next estimated *F*_IBD_ between all pairs of phased haplotypes. To define *F*_IBD_ between pairs of *isolates*, we took the maximum over the values for all combinations of haplotypes inferred from the isolates (**Figure 3C**). As expected, most pairs were effectively unrelated (median *F*_IBD_ <= 0.001, on the boundary of the parameter space), but a substantial fraction were related at the level of half-siblings or closer (*F*_IBD_ > 0.25, 4.0% of all pairs), including 1.3% of mainland-Zanzibar pairs.

**Figure 3.**
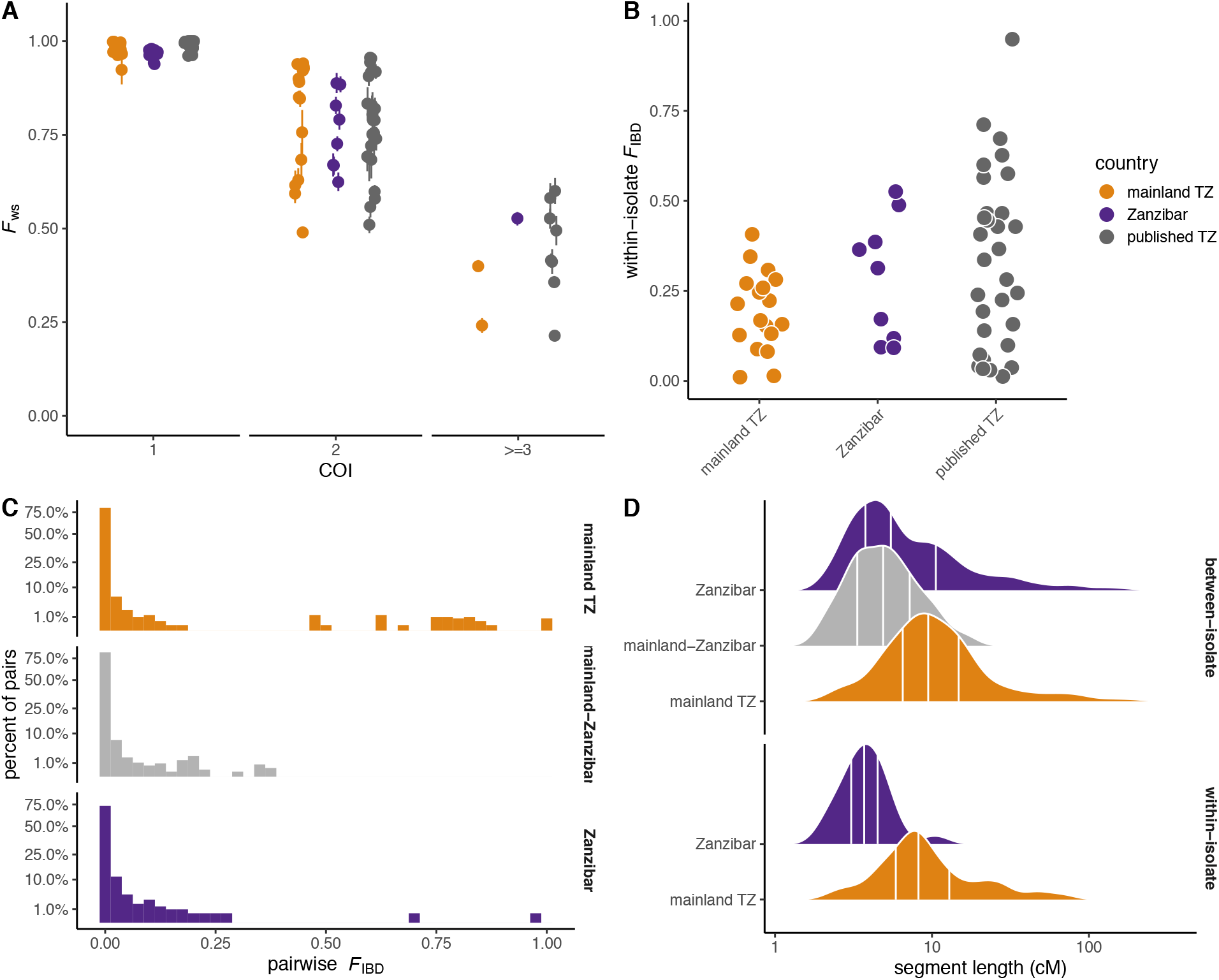
Complexity of infection and patterns of within- and between-host relatedness. (**A**) The *F*_ws_ index of within-host diversity, binned by complexity of infection (COI) estimated from genome-wide SNVs. Points colored by population. (**B**) Distribution of within-host relatedness, measured as the proportion of the genome shared IBD (*F*_IBD_) between strains, for isolates with COI > 1. Note that y-axis is on square-root scale. (**C**) Distribution of between-host relatedness, calculated from haplotype-level IBD. (**D**) Distribution of the length of segments shared IBD between (top) or within hosts (bottom). Segment lengths given in centimorgans (cM). Vertical lines mark 25th, 50th and 75th percentiles.

Long segments of the genome are shared IBD both within and between isolates. Mean within-isolate segment length was 5.7 cM (95% CI 4.1 -- 7.3 cM, *n* = 117) on the mainland and 3.7 cM (95% CI 2.8 -- 4.6 cM, *n* = 80) on Zanzibar in a linear mixed model with individual-level random effects; the full distributions are shown in **Figure 3D**. Segments shared between isolates within the mainland population (6.2 cM, 95% CI 5.9 -- 6.6 cM, *n* = 3279) were longer than segments shared within Zanzibar (4.5 cM, 95% 4.1 -- 4.8 cM, *n* = 592) or between mainland and Zanzibar populations (4.1 cM, 95% CI 3.9 -- 4.3 cM, *n* = 6506). After accounting for differences in segment length by population, difference in lengths of IBD segments detected between versus within individuals are not significant (mean difference −0.038 cM, 95% CI −0.10 -- 0.023 cM). In a random-mating population the length of a segment shared IBD between a pair of individuals with last common ancestor *G* generations in the past is exponentially-distributed with mean 100/(2\**G*) cM. The shared haplotypes that we observe, with length on the order of 5 cM, are thus consistent with shared ancestry in the past 10 generations -- although as many as half of such segments probably date back at least 20 generations [40]. In the presence of inbreeding, IBD sharing persists even longer in time.

Close relationships between isolates from the archipelago and the mainland suggest recent genetic exchange. We defined a threshold *F*_IBD_ > 0.25 because it implies that two isolates shared at least one common parent in the last outcrossing generation and therefore are related as recently as the last 1-2 transmission cycles, depending on background population dynamics. In principle this could result from importation of either insect vectors or human hosts. To investigate the latter possibility, we used a travel-history questionnaire completed by subjects from Zanzibar. Nine subjects reported travel to the mainland in the month before study enrollment; their destinations are shown in **Figure 4A**. We identified 10 pairs with *F*_IBD_ > 0.25 (marked by orange triangles in histogram in **Figure 4B**); all involved a single Zanzibar isolate from a patient who travelled to the coastal town of Mtwara (orange arc in **Figure 4A**). It is very likely that this individual represents an imported case. Overall, isolates from travelers had slightly higher mean pairwise relatedness to isolates from the mainland (mean *F*_IBD_ = 0.0020, 95% CI 0.0018 -- 0.0021) than did isolates from non-travellers (mean *F*_IBD_ = 0.0015, 95% CI 0.0014 -- 0.0016; Wilcoxon rank-sum test *p* = 1.8 × 10^−12^ for difference). But these relationships -- spanning 10 or more outcrossing generations -- are far too remote to be attributed to the period covered by the travel questionnaire. The pattern likely represents instead the presence of subtle population structure within Zanzibar.

**Figure 4.**
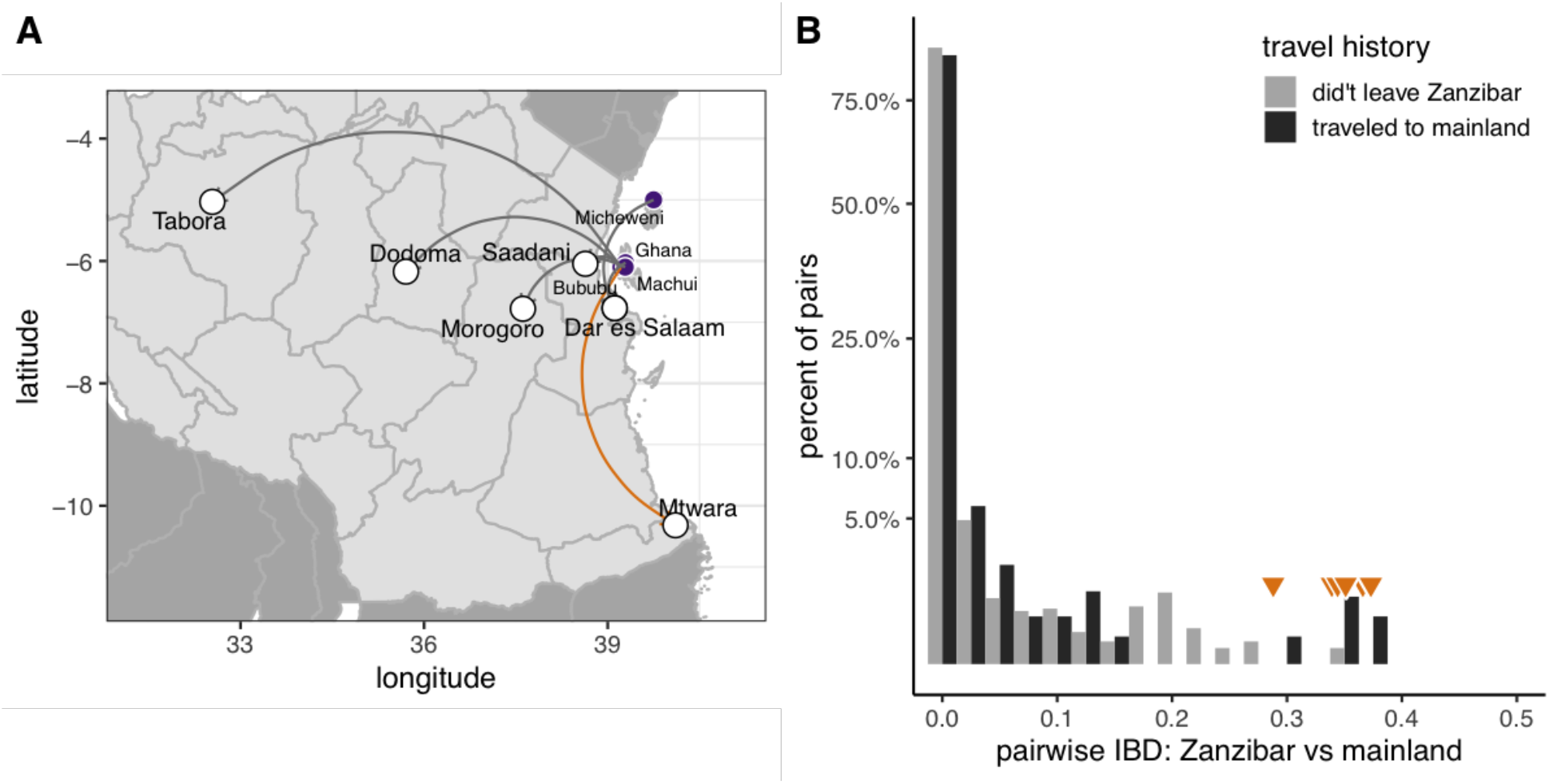
Travel history and parasite relatedness. (**A**) Reported destinations for 9 residents of Zanzibar who travelled to mainland Tanzania in the month before study enrollment. Orange arc shows destination of suspected imported case. (**B**) Pairwise IBD sharing between Zanzibar isolates from hosts with recent travel (dark bars) versus non-travelers (light bars). Values > 0.25 highlighted by orange triangles. Note that y-axis is on square-root scale.

### Demographic history of parasite populations

The distribution of IBD segment lengths carries information about the trajectory of effective population size in the recent past, up to a few hundred generations before the time of sampling. The site frequency spectrum and patterns of fine-scale linkage disequilibrium carry information about the more remote past. We used complementary methods to infer recent and remote population demography from phased haplotypes. First, we applied a non-parametric method [29] to infer recent effective population size (*N*_e_) from IBD segment lengths separately in mainland Tanzania and Zanzibar populations (**Figure 5A**). The method infers a gradual decline of several orders of magnitude in *N*_e_ over the past 100 generations to a nadir at *N*_e_ ~= 5,000 around 15-20 outcrossing generations before the time of sampling. Although the confidence intervals are wide, similar trajectories are inferred in all three populations.

**Figure 5.**
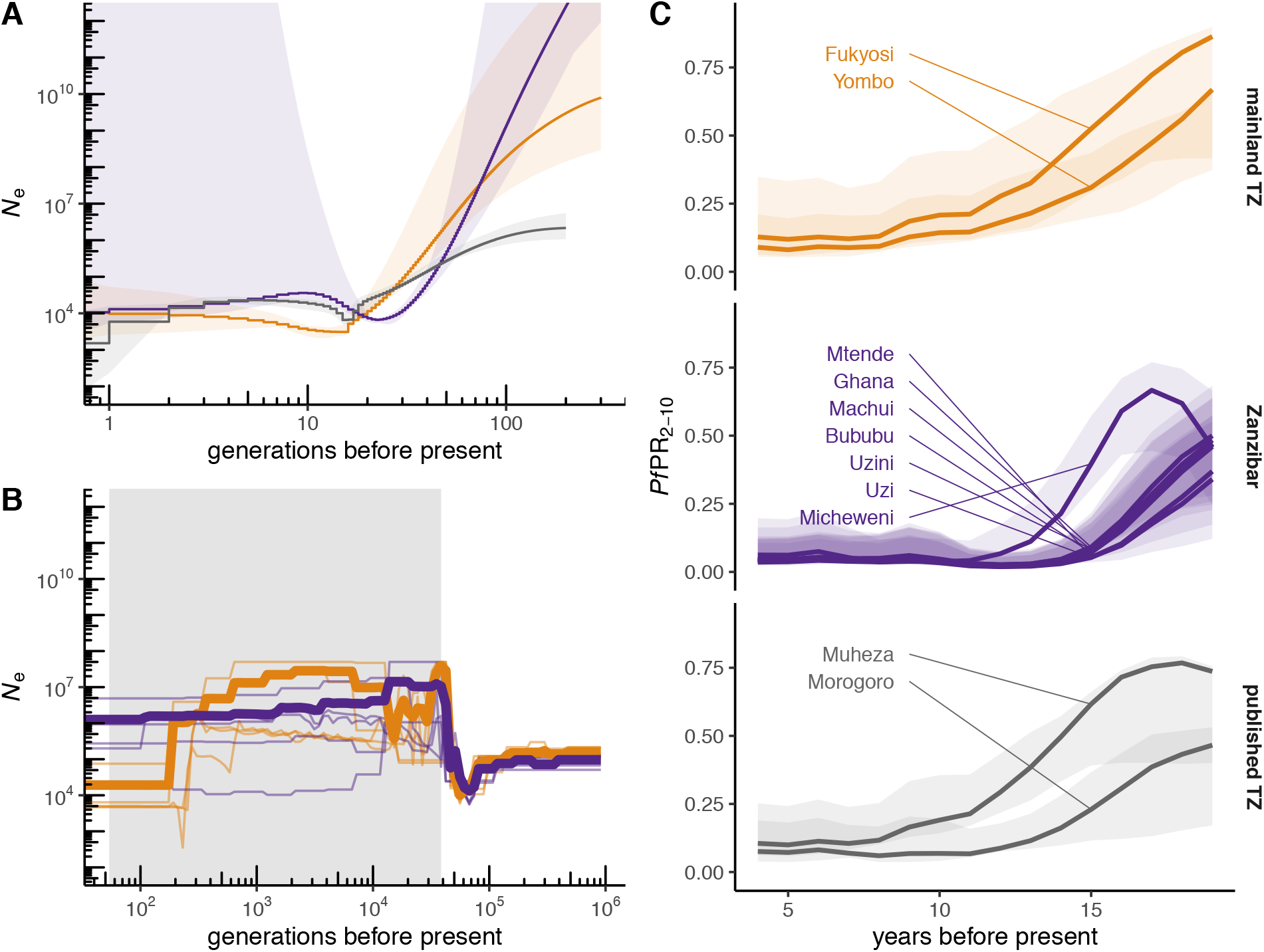
Comparison of historical parasite demography and infection prevalence. (**A**) Curves of recent historical effective population size (*N*_e_) reconstructed from IBD segments; shaded regions give 95% bootstrap CIs. (**B**) Effective population size in the more remote past, reconstructed from phased haplotypes. Thin lines, independent model runs; bold lines, model averages (see **Methods**). Shaded region, range of inferred split times between mainland and Zanzibar populations. Scale of y-axis matches panel A. (**C**) Estimated prevalence of *P. falciparum* infection from the Malaria Atlas Project at sampling sites for our cohorts (expressed as age-standardized prevalence rate among children aged 2-10 years, *Pf*PR_2-10_, in cross-sectional surveys); shaded regions give 95% credible intervals. Present = 2019.

Second, we inferred more remote population size histories jointly for mainland Tanzania and Zanzibar and attempted to estimate the split time between these populations using a sequentially Markovian coalescent method (Terhorst et al. 2017). This family of models has good resolution for relatively remote events but less precision in the recent past than models based on IBD segments. Our result (**Figure 5B**) supports a common ancestral population with *N*_e_ ~= 10^5^ individuals that underwent a sharp bottleneck followed by rapid growth around 50,000 generations before the present. The time at which the mainland and Zanzibar populations diverged could not be estimated precisely and may have been as recent as 50 or as ancient as 50,000 generations before the present. Trends in *N*_e_ were compared to local trends in parasite prevalence from the Malaria Atlas Project [30] (**Figure 5C**). Assuming an interval of approximately 12 months per outcrossing generation [41], the contraction in *N*_e_ may correspond in time to the decrease in prevalence brought about by infection-control measures over the past two decades.

### Natural selection and adaptation

Finally, we took several approaches to characterize the effects of natural selection on sequence variation in mainland and Zanzibar populations. The distribution of fitness effects (DFE) describes the relative proportion of new mutations that are deleterious, effectively neutral and beneficial and can be estimated from the frequency spectrum at putatively-neutral (synonymous) and putatively-selected (non-synonymous) sites (**Figure 6A**). Building on previous work in other organisms, we modeled the DFE in each population as a mixture of a gamma distribution (for deleterious mutations) and an exponential distribution (for beneficial mutations) [32]. We performed the inference using both the raw SFS and a smoothed representation of the SFS that is more numerically stable and found that results to be similar with both methods. Fitted parameter values are provided in **Supplementary Table 5** but the discretized representation of the DFE is more amenable to qualitative comparisons (**Figure 6B**).

**Figure 6.**
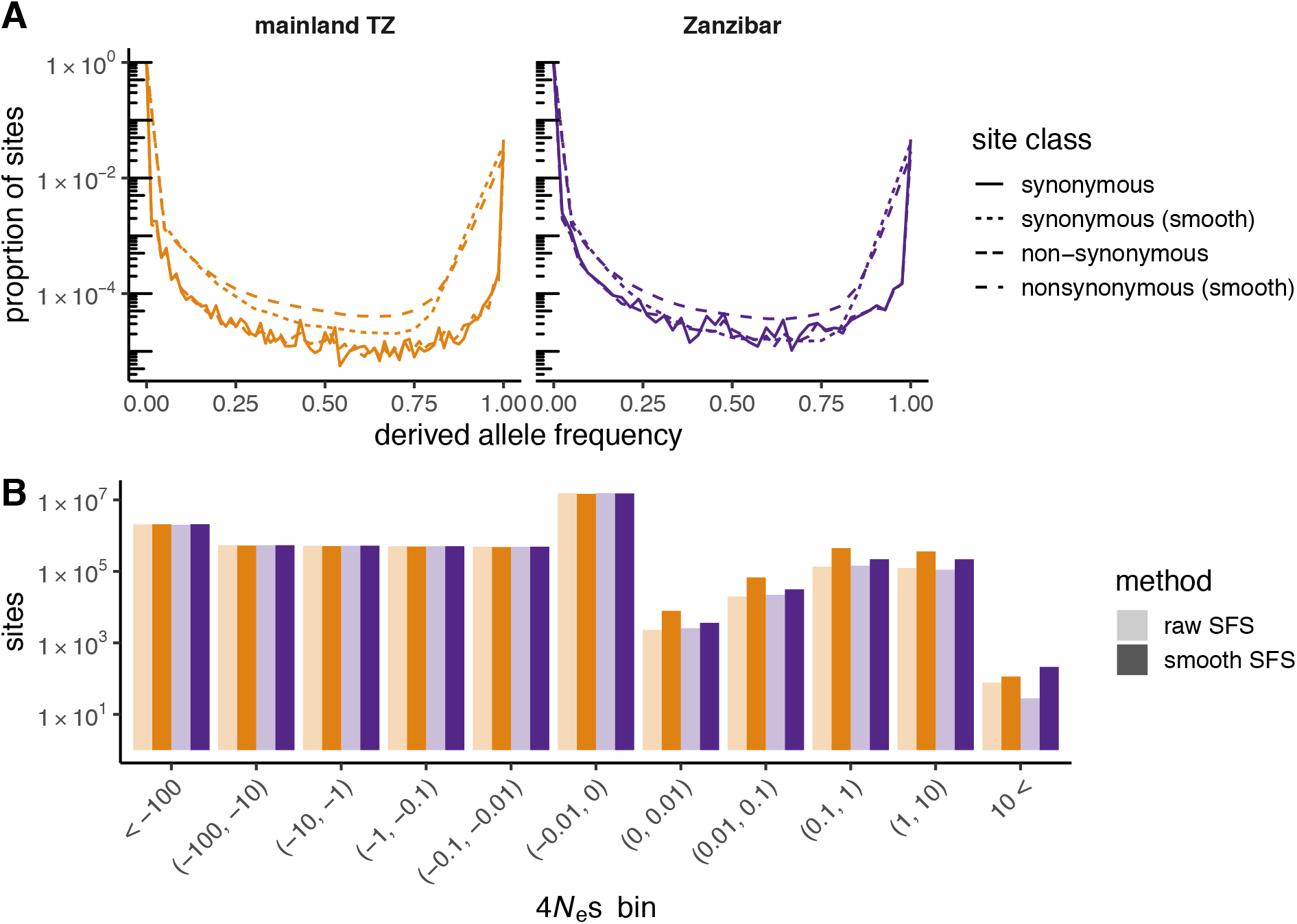
Characterizing the impact of natural selection on sequence variation. (**A**) Site-frequency spectra for putatively neutral (4-fold degenerate) and putatively-selected (0-fold degenerate) sites. (**B**) Inferred distribution of population-scaled selection coefficients (4*N*_e_*s*) for each population, shown in discrete bins. Dark bars, estimates from raw SFS; light bars, estimates from smoothed SFS. Note logarithmic scale for vertical axis in both panels.

The DFE allows us to estimate that 8.8% (mainland) and 5.2% (Zanzibar) of substitutions since the common ancestor with *P. reichenowi* have been fixed by positive selection; this quantity is known in some contexts as the “rate of adaptive evolution.” Differences in the DFE between populations are not statistically significant.The great majority of new mutations (mainland: 74%; Zanzibar: 76%) were expected to be very weakly deleterious (−0.01 < 4*N*_e_*s* < 0), and only a small minority were expected to be beneficial (4*N*_e_*s* > 0) (mainland: 4.5% [95% CI 2.7 -- 29%]; Zanzibar: 2.4% [95% CI 0.56 -- 50%]).

Although the DFE tells us the proportion of polymorphic sites under positive selection, it does not pinpoint which sites those are. To identify signals of recent, population-specific positive selection we used the XP-EHH statistic between mainland and Zanzibarian isolates [34]. Outliers in the XP-EHH scan, which we defined as standardized XP-EHH scores above the 99.9th percentile, represent candidates for local adaptation (**Supplementary Figure 2**). One-hundred four biallelic SNPs in 20 distinct genes passed this threshold (**Supplementary Table 6**). None of these have been associated with resistance to antimalarial drugs -- an important form of local adaptation in this species -- but one (PF3D7_0412300) has been identified in a previous selection scan [42]. Prevalences of 54 known drug-resistance loci are shown in **Supplementary Table 7** and is similar to previous reports in East Africa [43–45]. None of these mutations had *F*_st_ > 0.05 between mainland Tanzania and Zanzibar.

## DISCUSSION

Zanzibar has been the target of intensive malaria control interventions for nearly two decades following the early implementation of ACT therapies in 2003 [2]. Despite sustained vector control practices and broad access to rapid testing and effective treatment, malaria has not been eliminated from the archipelago [2]. Here we use WGS of *P. falciparum* isolates from Zanzibar and nearby sites on the mainland to investigate ancestry, population structure and transmission in local parasite populations. Our data place Tanzanian parasites in a group of east African populations with broadly similar ancestry and level of sequence diversity. We find minimal signal of differentiation between mainland and Zanzibar isolates.

The most parsimonious explanation for our data is a source-sink scenario, similar to a previous report in Namibia [46], in which importation of malaria from a region of high but heterogeneous transmission (the mainland) is inhibiting malaria elimination in a pre-elimination area (Zanzibar). Using WGS we show that the parasite population on the islands remains genetically almost indistinguishable from regions on the mainland of Tanzania. We can identify numerous long segments of the chromosomes that are shared between the populations, on the order of 5 cM, suggesting that genetic exchange between the populations has occurred within the last 10-20 sexual generations. In addition, we identify a Zanzibar isolate that is related at the half-sibling level to a group of mutually-related mainland isolates. This likely represents an imported case and provides direct evidence for recent, and likely ongoing, genetic exchange between the archipelago and the mainland. These observations suggest that parasite movement from the mainland to the archipelago is appreciable and may be a significant hurdle to reaching elimination.

Human migration is critical in the spread of malaria [47], thus the most likely source for importation of parasites into Zanzibar is through human travel to high-risk malaria regions. There have been multiple studies on the travel patterns of Zanzibarian residents as it relates to importation of malaria [48–50], one of which estimated that there are 1.6 incoming infections per 1,000 inhabitants per year. This is also in accordance with the estimate of about 1.5 imported new infections out of a total of 8 per 1000 inhabitants in the recent epidemiological study [2]. None of these studies have leveraged parasite population genetics to understand importation patterns. Though our study is small, our data suggests that genetics can potentially provide additional insight into the impacts of travel and the corridors of parasite migration to Zanzibar.

Malarial infections in Africa are highly polyclonal. This within-host diversity poses technical challenges but also provides information on transmission dynamics. Approximately half of isolates from both the mainland and Zanzibar represent mixed infections (COI > 1), similar to estimates in Malawian parasites with similar ancestry [15]. We found that a widely-used heuristic index (*F*_ws_) is qualitatively consistent with COI estimated by haplotype deconvolution [51], but has limited discriminatory power in the presence of related lineages in the same host. Furthermore, median within-host relatedness (*F*_IBD_) is ~0.25, the expected level for half-siblings, in both mainland and Zanzibar populations. This strongly suggests frequent co-transmission of related parasites in both populations [39]. Our estimates of *F*_IBD_ are within the range of estimates from other African populations and add to growing evidence that mixed infections may be predominantly due to co-transmission rather than superinfection even in high-transmission settings [52,53].

Intensive malaria surveillance over the past several decades provides an opportunity to compare observed epidemiological trends to parasite demographic histories estimated from contemporary genetic data. Our estimates of historical effective population size (*N*_e_) support an ancestral population of approximately 10^5^ individuals that grew rapidly around 10^4^ generations ago, then underwent sharp contraction within the past 100 generations to a nadir around 10-20 generations before the present. We were unable to obtain stable estimates of the split time between the mainland and Zanzibar populations, either with a coalescent-based method (**Figure 5B**) or with method based on the diffusion approximation to the Wright-Fisher process (not shown) (Gutenkunst et al. 2009). This is not surprising given that the shape of joint site frequency spectrum (**Supplementary Figure 3**), summarized in low *F*_st_ genome-wide, is consistent with near-panmixia. The timing and strength of the recent bottleneck appears similar in our mainland Tanzania and Zanzibar isolates and coincides with a decline in the prevalence of parasitemia. However, we caution that the relationship between genetic and census population size -- for which prevalence is a proxy -- is complex, and other explanations may exist for the observed trends.

Finally, we make the first estimates of the distribution of fitness effects (DFE) in *P. falciparum*. Although the impact of selection on genetic diversity in this species has long been of interest in the field, previous work has tended to focus on positive selection associated with resistance to disease-control interventions. The DFE is a more fundamental construct that has wide-ranging consequences for the evolutionary trajectory of a population and the genetic architecture of phenotypic variation [54]. We find that the overwhelming majority of new alleles are expected to be deleterious (*N*_e_*s* < 0) but most (~75%) have sufficiently small selection coefficients that their fate will be governed by drift. The proportion of new mutations expected to be beneficial -- the “target size” for adaption-- is small, on the order 1-2%. Together these observations imply that even in the presence of ongoing human interventions, patterns of genetic variation in the Tanzanian parasite population are largely the result of drift and purifying selection rather than positive selection. We note that these conclusions are based on the core genome and may not hold for hypervariable loci thought to be under strong selection such as erythrocyte surface antigens. Furthermore, the complex lifecycle of *Plasmodium* species also departs in important ways from the assumptions of classical population-genetic models [55]. The qualitative impact of these departures our conclusions is hard to determine.

## CONCLUSION

The elimination of malaria from Zanzibar has been a goal for many years. Here we present genomic evidence of continued recent importation of *P. falciparum* from mainland Tanzania to the archipelago. Reducing this importation is likely to be an important component of reaching the elimination end game. Investigation of methods to do this, such as screening of travelers or mass drug treatment, is needed. However, the high degree of connectivity between the mainland and the Zanzibar archipelago will make this challenging. We are encouraged by evidence that parasite populations in the region are contracting (**Figure 5**). These declines are likely due to decreasing transmission but need to be interpreted with caution, as they may also be due to other factors that impact effective population size estimates, including violation of model assumptions. The data suggests that larger studies of the relationship between Zanzibarian and mainland parasites will enable further more precise estimates of corridors of importation based on parasite genetics. Genomic epidemiology has the potential to supplement traditional epidemiologic studies in Zanzibar and to aid efforts to achieve malaria elimination on the archipelago.

## ETHICAL APPROVALS AND CONSENT TO PARTICIPATE

This analysis was approved by the IRBs at the University of North Carolina at Chapel Hill, Muhimbili University of Health and Allied Sciences (MUHAS), Zanzibar Medical Research Ethical Committee and the Regional Ethics Review Board, Stockholm, Sweden.

## CONSENT FOR PUBLICATION

Not applicable.

## AVAILABILITY OF DATA AND MATERIAL

Sequencing reads were deposited into the NCBI SRA (Accession numbers: pending). Code is available through GitHub (https://github.com/IDEELResearch). This publication uses data from the MalariaGEN *P. falciparum* Community Project (www.malariagen.net/projects/p-falciparum-community-project) as described in [35]. Genome sequencing was performed by the Wellcome Trust Sanger Institute and the Community Projects is coordinated by the MalariaGEN Resource Centre with funding from the Wellcome Trust (098051, 090770). This publication uses data generated by the Pf3k project (www.malariagen.net/pf3k) which became open access in September 2016.

## AUTHOR CONTRIBUTIONS

APM, NFB and JBP designed experiments, conducted analysis and wrote the manuscript. BN, EL, MM, and UM collected samples and participated in manuscript preparation. MD conducted laboratory work and participated in manuscript preparation. DLF helped develop software and participated in manuscript preparation. JAB, AM, AB and JJJ helped conceive the study, contributed to the experimental design and wrote the manuscript.

## ACKNOWLEDGEMENTS

We would like to thank the communities and participants who took part in these studies. We would also like to thank Molly Deutsch-Feldman for helping to optimize the sWGA protocol.

