## Supplementary text, tables and figures for "Falciparum malaria from coastal Tanzania and Zanzibar remains highly connected despite effective control efforts on the archipelago"

This file includes supplemental material for:

### Contents

| | |
|---|---|
| Supplemental Materials | Page 2 |
| Supplemental Tables | Page 5 |
| Supplemental Table 1 | Page 5 |
| Supplemental Table 2 | Page 6 |
| Supplemental Table 3 | Page 7 |
| Supplemental Table 4 | Page 8 |
| Supplemental Table 5 | Page 9 |
| Supplemental Table 6 | Page 10 |
| Supplemental Table 7 | Page 11 |
| Supplemental Table 8 | Page 12 |
| Supplemental Figures | Page 13 |
| Supplemental Figure 1 | Page 13 |
| Supplemental Figure 2 | Page 14 |
| Supplemental Figure 3 | Page 15 |
| Supplemental Figure 4 | Page 16 |
| Supplemental References | Page 17 |

### SUPPLEMENTAL MATERIALS

#### *Design of primers for selective whole genome amplification (sWGA)*

We developed sWGA primer sets using ‘swga’, a recently developed program that supports sWGA primer design by identifying and ranking candidate primer sets(Clarke et al. 2017). These sets are designed to selectively amplify a “foreground” genome (e.g. *P. falciparum*), with minimal amplification of a “background” genome (e.g. human), and hard limits on primer binding to “exclusion” genomes (e.g. human mitochondria).

We used the following genomes during primer design:

1) Foreground: *P. falciparum* 3D7 (assembly version 3.0, PlasmoDB version 13.0)
2) Background: Human (GRCH38.p7)
3) Exclusion: Human mitochondrial DNA (NC_012920.1, included in GRCH38.p7 above)

We restricted candidate primers to those with annealing temperatures ranging from 18-30C and permitted a maximum of 14 primers per primer set. Otherwise, default parameters were used to identify candidate sets consisting of 5-12mer primers. We ran the ‘swga’ program in parallel on the University of North Carolina Killdevil computing cluster, with one run completely excluding primers that bound the human mitochondrial genome and a second run that permitted a maximum of three primer binding sites. We allowed each to run for 4 days using a dedicated node with 12 processors, producing approximately 200,000 candidate primer sets that passed all filters.

We then chose candidate primer sets for testing in the laboratory. First, we selected 6 candidate sets based on the ‘swga’ default scoring algorithm, which incorporates the mean distance between primer binding sites on the foreground genome (target primer binding site density), the foreground binding Gini coefficient (a measure of evenness of primer binding sites across the foreground genome), and the background binding distance (background primer binding site density) as follows:

$$\frac{(Mean\ foreground\ binding\ distance) \times (Foreground\ distance\ Gini\ coefficient)}{(Mean\ background\ binding\ distance)}$$

Second, we selected 5 candidate sets with the lowest mean foreground binding distance not already selected based on their score, including 3 with no predicted human mitochondrial binding and 2 with ≤3 human mitochondrial binding sites. Characteristics of candidate primer sets are provided in **Supplemental Table 8**.

#### *Initial testing of candidate sWGA primer sets*

Testing of candidate primer sets involved sWGA performed as previously described(Sundararaman et al. 2016). A 5% mixture of *P. falciparum* 3D7 strain DNA in human DNA with a total concentration of 1ng/uL was used as starting template. In brief, starting template was first equilibrated at 35C for 5 minutes. For each primer set, we prepared a 50uL reaction mixture as follows: 3.5uM of pooled primers, 1x *phi29* reaction buffer (New England Bio Labs), 1mM of deoxynucleotide solution (New England Bio Labs), 10 ug of bovine serum albumin, 30 units of *phi29* polymerase (New England Bio Labs), nuclease-free water, and 10ng of starting DNA template. Next, sWGA was performed in a thermocycler using the following step-down protocol: 35C for 1 hour, 34C for 1 hour, 33C for 1 hour, 32C for 1 hour, 31C for 1 hour, 30C for 16 hours, and 65C for 10 minutes. Amplification product was cleaned up using Agencourt AmpureXP beads (Beckman Coulter). For comparison, we also performed two-step sWGA using primer sets 6A and 8A designed and reported by Sundararaman *et al(Sundararaman et al. 2016).*

We performed an initial assessment of enrichment by primer set using duplex, quantitative PCR (qPCR) targeting the single-copy *P. falciparum* lactate dehydrogenase (*pfldh*) gene and the human beta tubulin (HumTuBB) gene as previously described(Mwandagalirwa et al. 2017). We calculated enrichment by comparing Ct values of sWGA product to those of unamplified starting template. This approach provides an imperfect estimate of enrichment, as breadth of coverage after sWGA is often uneven and involves “jackpotting” of specific regions, but allowed us to focus sequencing efforts on a smaller group of primer sets. Results of qPCR and initial enrichment estimates are provided in **Supplemental Table 8.**

***Whole-genome sequencing using candidate primer sets***

We selected sWGA product from reactions performed with primer sets JP1, JP5, JP6, JP7, JP9, and JP11 for sequencing. These primer sets were drawn from all four categories of primer set designs (mitochondrial binding permitted *vs.* not permitted, ranking by score *vs.* foreground binding distance), prioritizing unique sets with the greatest enrichment as estimated by qPCR. Cleaned product was acoustically sheared using a Covaris E220 machine and prepared for sequencing using a Kappa Hyper prep kit. Libraries were pooled to achieve an equimolar mixture and sequenced on a single MiSeq run, using 300bp paired-end chemistry. Paired-end reads were aligned to the *P. falciparum* 3D7 reference genome (assembly version 3, PlasmoDB version 13) with `bwa mem`. In order to estimate enrichment, we also performed a separate alignment to the human GRCH38.p7 reference genome using `bwa mem`.

We calculated the percent of total reads per library that mapped to the *P. falciparum* and human genomes, respectively, as a measure of enrichment. Compared to the unamplified 5% *P. falciparum* (95% human) starting template, total enrichment ranged from 7.6- to 15.5-fold. We also calculated the median *P. falciparum* coverage (≥1x) by chromosome as a measure of sequencing breadth across the target genome. Median coverage ranged from 68.4% to 99.7%. Detailed results by primer set are provided in the **Supplemental Table 8 and Supplementary Figure 3.** Primer set JP9 achieved the best results when both enrichment (14.4-fold) and breadth (99.6%) were taken into consideration.

***Characterization of the best performing sWGA primer set*** We further characterized the performance of the JP9 primer set, in addition to the JP11 set, and the two-step 06A+08A sWGA approach previously described by Sundararaman *et al.* using serial dilutions of *P. falciparum* DNA in a background of human DNA. We performed each of these sWGA reactions in singleton using 1% *P. falciparum* 3D7 DNA (99% human DNA) with a total DNA concentration of 1ng/uL. We also performed each sWGA reaction in triplicate using a mixture of DNA from two *P. falciparum* strains (50% 3D7, 20% FCR3, and 30% DD2) serially diluted in human DNA to achieve 0.1%, 0.01%, and 0.001% *P. falciparum* DNA. sWGA product was cleaned up, sheared, and library prepped as described above. An equimolar pool of all libraries was sequenced on a single HiSeq 2500 lane, using 150bp, paired-end chemistry. Raw reads were aligned to the *P. falciparum* 3D7 genome (version 3.0, PlasmoDB version 13.0) using `bwa mem`. Duplicates were marked using the Picard tool `MarkDuplicates`. Indels were identified and realigned using GATK's `RealignerTargetCreator` and `IndelRealigner` tools, respectively.

We observed evidence of enrichment in all replicates containing ≥0.01% *P. falciparum* DNA, a finding generally in line with that reported by Oyola *et al* (**Supplemental Figure 4**)(Oyola et al. 2016). While all tested sWGA primer sets demonstrated evidence of enrichment, the JP9 set afforded superior breadth of coverage compared to JP11 and improved workflow compared to the published, two-step assay. We also assessed a two-step sWGA approach involving amplification with the JP9 set followed by amplification with the JP11 set but did not observe improved enrichment or breadth of coverage (data not shown).

***Revised sWGA protocol***

For clinical DBS samples, we revised our original sWGA approach in order to incorporate recently published methods described by Oyola *et al.* and Clarke *et al.* into our workflow. Specifically, we increased the concentration of the individual sWGA primers and the amount of total DNA input in each reaction.

In brief, we prepared a 50uL reaction mixture on ice, consisting of 2.5uM of each individual primer, 1x *phi29* reaction buffer (New England Bio Labs), 1mM of deoxynucleotide solution (New England Bio Labs), 10 ug of bovine serum albumin, 30 units of *phi29* polymerase (New England Bio Labs), nuclease-free water, and 40ng of starting DNA template. sWGA was performed in a thermocycler using the following step-down protocol: 35C for 5 min, 34C for 10 min, 33C for 15 min, 32C for 20 min, 31C for 30 min, 30C for 16 hours, and 65C for 15 minutes. Amplification product was cleaned up using Agencourt AmpureXP beads (Beckman Coulter).

### SUPPLEMENTAL TABLES

**Supplemental Table 1 - Clinical characteristics of cohorts.** Common baseline characteristics among the three cohorts considered in this study. Overall, samples that did not pass quality-control (QC) metrics were not significantly different to those that were included in the study with the exception of the Zanzibar, Asymptomatic cohort, where low parasitemia samples were preferentially lost. Given the low number of samples, clinical characteristics were compared using Fisher's exact test for categorical variables and Wilcoxon rank sum test was used for continuous variables. Age was reported in years, while parasitemia was reported in parasites per microliter.

| Study | Covariates | Failed QC | Passed QC | P-value |
|---|---|---|---|---|
| Zanzibar, Symptomatic | n | 12 | 17 | |
| Zanzibar, Symptomatic | Age (median [IQR]) | 11.50 [6.75, 15.75] | 13.00 [4.00, 23.00] | 0.706 |
| Zanzibar, Symptomatic | Male (%) | 10 (83.3) | 10 (58.8) | 0.234 |
| Zanzibar, Symptomatic | Parasitemia (median [IQR]) | 44200.00 [31790.00, 62950.00] | 57080.00 [40800.00, 100360.00] | 0.33 |
| Zanzibar, Asymptomatic | n | 30 | 4 | |
| Zanzibar, Asymptomatic | Age (median [IQR]) | 20.00 [16.00, 31.50] | 12.00 [11.25, 24.00] | 0.349 |
| Zanzibar, Asymptomatic | Male (%) | 16 (53.3) | 3 (75.0) | 0.613 |
| Zanzibar, Asymptomatic | Parasitemia (median [IQR]) | 126.00 [72.36, 405.93] | 31940.57 [18463.98, 56426.16] | <0.001 |
| Bagamoyo, Symptomatic | n | 7 | 36 | |
| Bagamoyo, Symptomatic | Age (median [IQR]) | 8.00 [6.25, 9.75] | 7.50 [5.25, 9.00] | 0.744 |
| Bagamoyo, Symptomatic | Male (%) | 4 (57.1) | 21 (58.3) | 1 |
| Bagamoyo, Symptomatic | Parasitemia (median [IQR]) | 1890.00 [512.00, 3850.00] | 1563.50 [582.75, 2612.50] | 0.984 |

**Supplemental Table 2 - Primer sequences for the custom, *P. falciparum*-specific selective whole genome amplification (sWGA) reaction (set JP9)**

| Primer Name | Primer Sequence |
|---|---|
| JP9a | ATATAATA*A*C |
| JP9b | TAATATAAT*A*A |
| JP9c | TATAATAA*G*A |
| JP9d | TATAGTAA*T*A |
| JP9e | TATATGAT*A*A |
| JP9f | TATCATAA*T*A |
| JP9g | TATGATAA*T*A |
| JP9h | TCGTAA*T*A |

* Denotes phosphorothioate bonds.

**Supplemental Table 3 - Publically available genomes used in analysis.**

Table submitted as separate file due to size.

**Supplemental Table 4 - Complexity of infection (COI)**

| **Population** | **COI = 1** | **COI = 2** | **COI >= 3** | **Total** |
|---|---:|---:|---:|---:|
| Mainland TZ | 18 | 16 | 2 | 36 |
| Zanzibar | 12 | 8 | 1 | 21 |
| **Total** | 30 | 24 | 3 | 57 |

**Supplemental Table 5 - Fitted parameters for distribution of fitness effects (DFE) analysis.**

| Population | Parameter | Estimate | 95% boostrap CI |
|---|---|---|---|
| Mainland TZ | $p_b$ | 0.045 | 0.027 -- 0.29 |
| | *alpha* | 0.089 | 0.046 -- 0.59 |
| | $-S_d$ | 11210 | 4736 -- 11633 |
| | *m* | 0.0137 | 0.0118 -- 0.0138 |
| | $S_b$ | 1.1 | 0.78 -- 1.4 |
| Zanzibar | $p_b$ | 0.024 | 0.0056 -- 0.50 |
| | *alpha* | 0.052 | 0.0090 -- 1.0 |
| | $-S_d$ | 9570 | 0.0002 -- 1262 |
| | *m* | 0.0138 | 0.0107 -- 0.208 |
| | $S_b$ | 1.3 | 0.21 -- 20 |

Notation for parameters is adapted from the `polyDFE` manual (Tataru et al. 2017): $p_b$, fraction of new alleles that are beneficial; $-S_d$, mean population-scaled selection coefficient for deleterious mutations; *m*, shape parameter for gamma (deleterious) component of DFE; $S_b$, mean population-scaled selection coefficient for beneficial mutations; *alpha*, proportion of substitutions fixed by natural selection (ie. rate of adaptive evolution.) Values correspond to analysis with smoothed SFS.

**Supplemental Table 6 - Significant genes identified by XP-EHH analysis between mainland and Zanzabarian parasite populations.**

For each of the biallelic SNPs identified, the genomic position (CHROM, POS) and centimorgan (cM) position is provided. The description column was taken directly from the *P. falciparum* 3D7 GFF3 file (PlasmoDB version 38: https://plasmodb.org/common/downloads/release-38/Pfalciparum3D7/gff/data/PlasmoDB-38_Pfalciparum3D7.gff). The remaining columns were downloaded from PlasmoDB, where column descriptions can be found (https://plasmodb.org/), using `Rselenium`. The ontology column represent the ontology term associated with the gene-ontology ID, the GO Slim ID column represents a subset of GO terms that are used by EuPathDB, and the GO Slim Term Name is the property or associated activity of the GO Slim ID that has been identified.

Table submitted as separate file due to size.

**Supplemental Table 7 - Drug resistance allele prevalence:** The prevalence of putative drug-resistance loci observed among the Tanzania and Zanzibar samples. Prevalence was calculated as the total number of samples with evidence of a putative drug-resistance SNP (i.e. either a homozygous putative-resistance or heterozygous genotype call) divided by the total number of samples with a genotype call at that loci. Of note, the *crt* K75E mutation genotyped as a complex multinucleotide variant in our cohort and was excluded from this analysis.

Table submitted as separate file due to size.

**Supplemental Table 8 - Characteristics of candidate sWGA primer sets.** A) Candidate primer sets by category. B) Initial assessment of candidate sWGA primer sets using *pfldh* qPCR as a crude measure of enrichment and selection of sets for testing by NGS. C) Results from a single Illumina MiSeq run of pooled sWGA product from the best performing primer sets.

Table submitted as separate file due to size.

### SUPPLEMENTAL FIGURES

**Supplemental Figure 1 - Mutational spectrum for SNVs.** Density of SNVs per kb of sequence in the core genome, normalized for base composition. Classes of mutations are collapsed by strand symmetry; ":" denotes base-pairing, eg. "A:T>C:G" is all A>C or T>G changes. Transitions, open circles; transversions, filled circles.

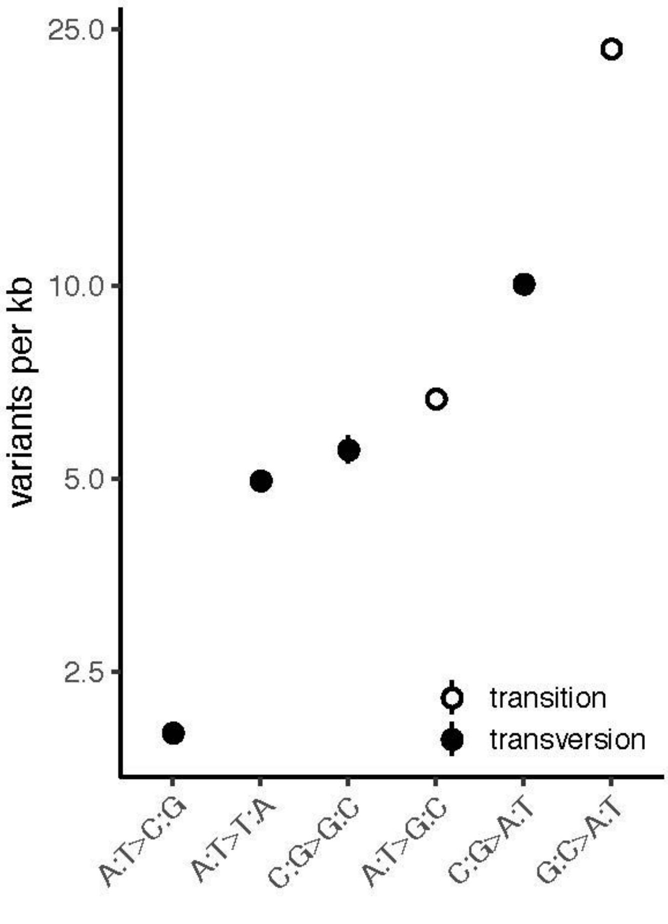

**Supplemental Figure 2 - XP-EHH genome scan**. Each point shows standardized cross-population extended haplotype homozygosity (XP-EHH) score (upper panel) or the associated *p*-value (lower panel) for 103,982 biallelic SNVs across the core genome. The populations compared are all monoclonal isolates from Tanzania (*n* = 18) and Zanzibar (*n* = 12). Outlier sites (-log10 p-value >= 99.9th percentile, red dashed line) are marked in red. Position of known drug-resistance loci marked in dark blue.

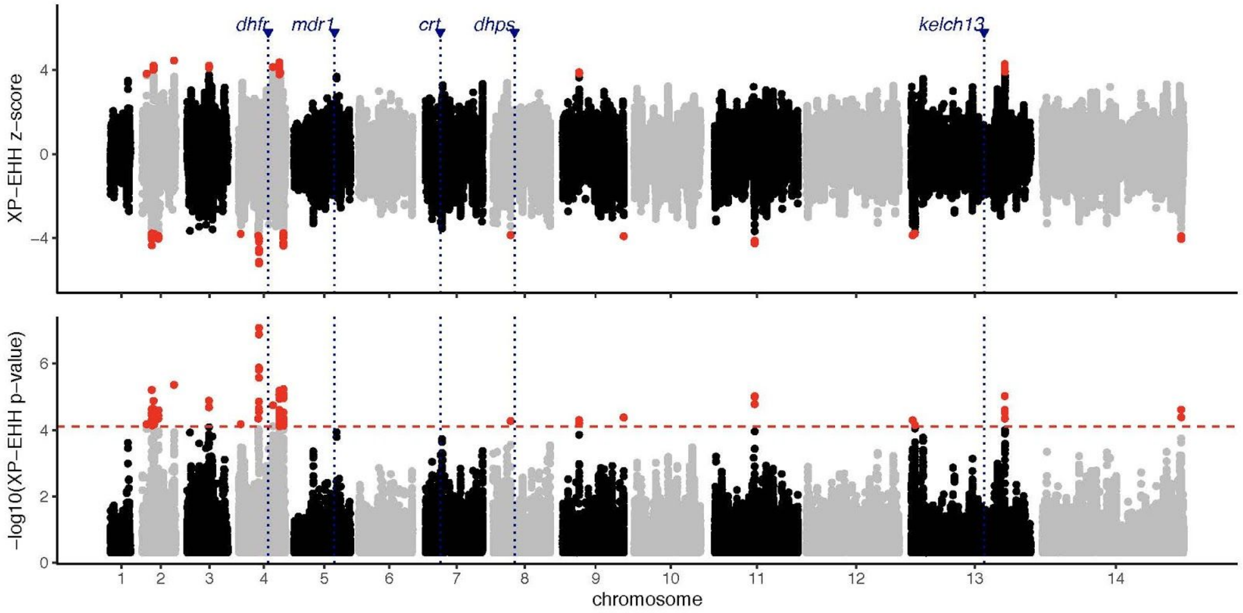

**Supplemental Figure 3 - Coverage of candidate sWGA primer sets.** The proportion of the genome covered with X-fold depth for each respective sWGA primer-set.

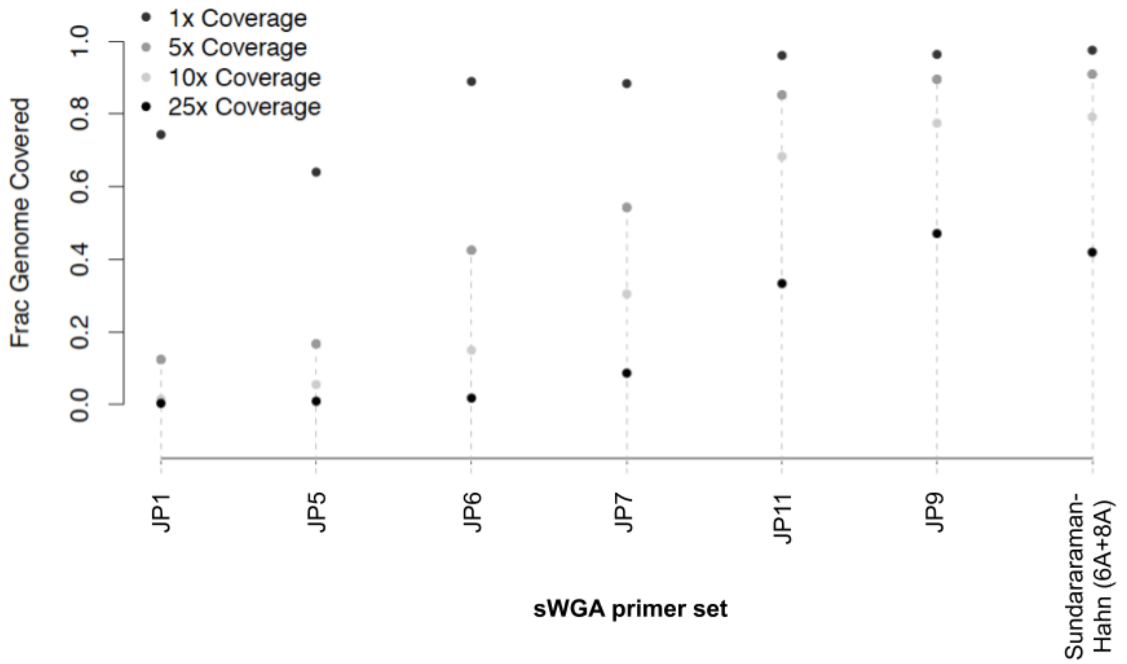

**Supplemental Figure 4 - Coverage of sWGA primers on known mixtures.** The proportion of the genome covered with X-fold depth for each respective sWGA primer-set. *P. falciparum* genomic DNA was diluted to 1% ("X_1"), 0.1% ("X_01"), and 0.01% ("X_001") with human genomic DNA for each mixture. Mono-infections with 1% *P. falciparum* 3D7 ("mono") were also considered. Finally, two mixtures that did not undergo sWGA with *P. falciparum* 3D7 DNA only (i.e. no human DNA; "MixUnampNoHuman") and 1% *P. falciparum* 3D7 DNA ("MixUnamp") were sequenced as template controls.

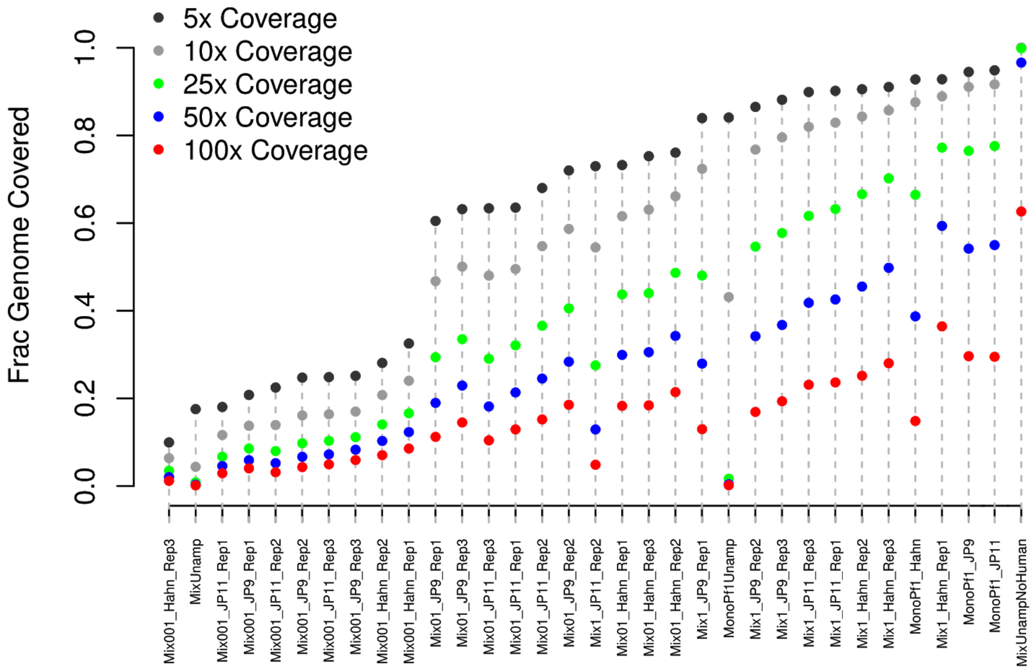
