## Supplementary figures and images for "Falciparum malaria from coastal Tanzania and Zanzibar remains highly connected despite effective control efforts on the archipelago"

### Figure S1

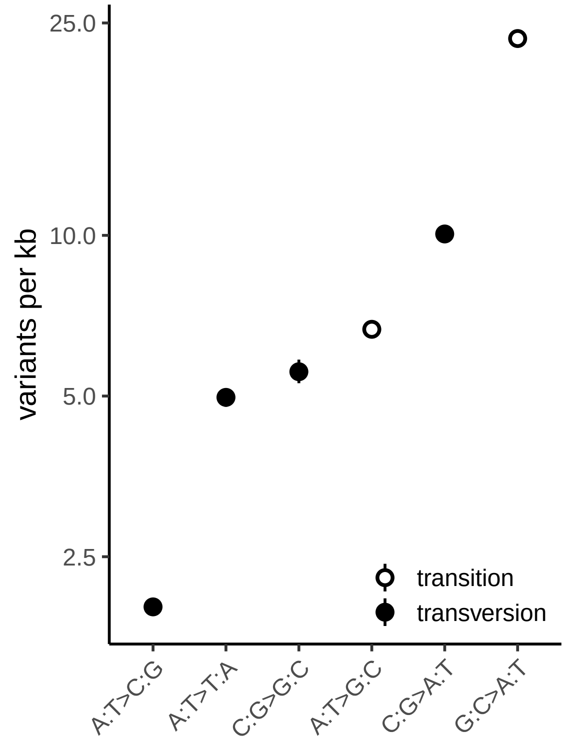

### Figure S2

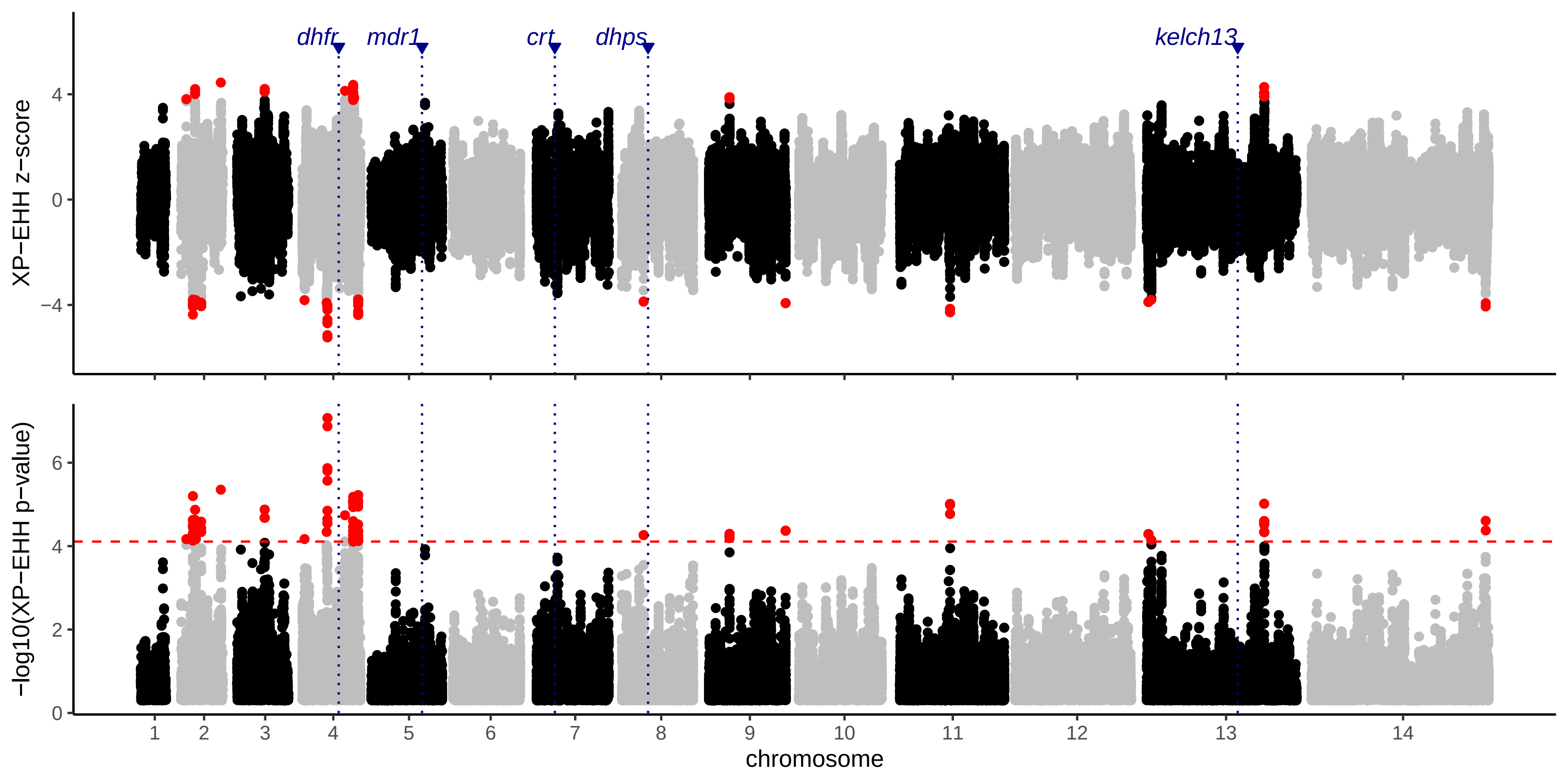

### Figure S3

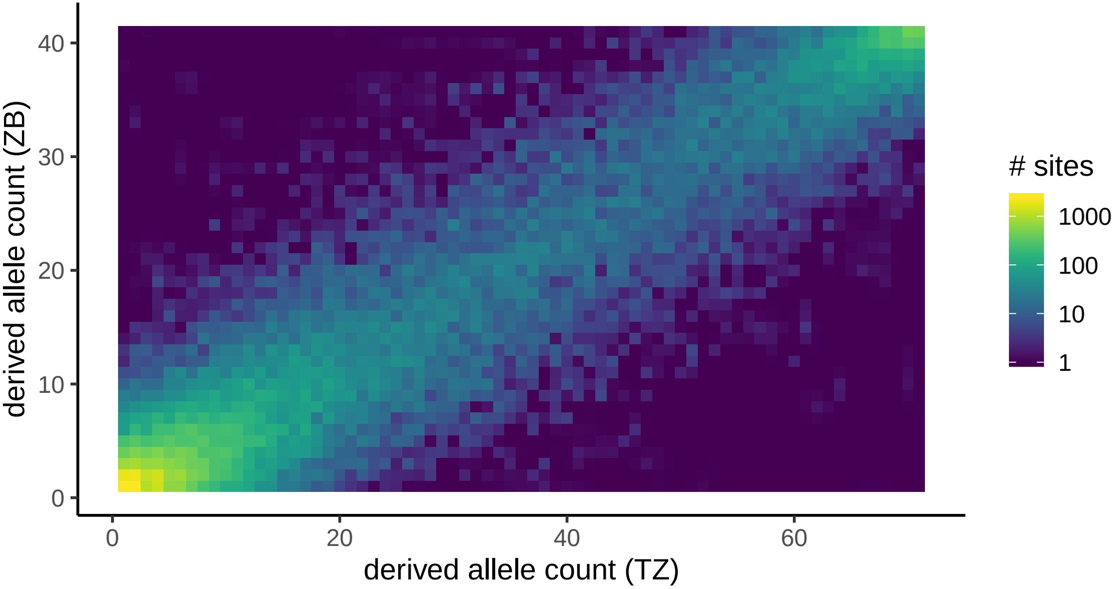
